# GatorAffinity: Boosting Protein-Ligand Binding Affinity Prediction with Large-Scale Synthetic Structural Data

**DOI:** 10.1101/2025.09.29.679384

**Authors:** Jinhang Wei, Yupu Zhang, Peter A Ramdhan, Zihang Huang, Gustavo Seabra, Zhe Jiang, Chenglong Li, Yanjun Li

## Abstract

Protein–ligand binding affinity prediction is a fundamental task in computational drug discovery. Although substantial efforts have been made to enhance prediction accuracy using data-driven approaches, progress remains limited by persistent data scarcity. The widely used PDBbind dataset, for example, contains fewer than 20, 000 experimental structures with annotated binding affinities, while a vast number of affinity measurements remain underutilized due to missing structural data. Here, we investigate this untapped potential by curating more than 450, 000 synthetic protein–ligand complexes annotated with *K*_*d*_ and *K*_*i*_ values using the Boltz-1 structure prediction model. Building on this unprecedented scale of synthetic data, further augmented with over 1 million synthetic complexes from the recently released SAIR database annotated with *IC*_50_ values, we develop GatorAffinity, a geometric deep learning–based scoring function pretrained on large-scale synthetic data and fine-tuned using high-quality experimental structures from PDBbind. Extensive evaluation on a leak-proof benchmark demonstrates that GatorAffinity significantly outperforms state-of-the-art affinity prediction methods, offering superior accuracy and generalizability. Our findings show that augmenting available experimental data with synthetic complexes can effectively address the data scarcity challenge while maintaining strong predictive reliability. By releasing the pretrained GatorAffinity model and the large-scale synthetic dataset GatorAffinity-DB, we provide a scalable and reproducible foundation for affinity prediction, virtual screening, and broader structure-based drug design applications (https://github.com/AIDD-LiLab/GatorAffinity).

## 1 Introduction

Drug discovery is a complex and highly challenging process. A fundamental objective is to identify small molecules that can bind specifically to protein targets and modulate their biological functions, a strategy known as structure-based drug design (SBDD). Traditional methods are highly dependent on laboratory techniques, including high-throughput screening (HTS) and structural biology techniques (such as X-ray crystallography and nuclear magnetic resonance), all of which are significantly labor-intensive and require substantial time and economic costs [1]. Therefore, accurately predicting protein–ligand binding affinity has become a critical step for efficiently exploring the vast chemical space, reducing costs, and accelerating drug development.

Classical methods for affinity prediction typically rely on fixed, theory-driven functional forms applied to hand-crafted features derived from protein–ligand complexes. Although some approaches, such as free energy perturbation [2], thermodynamic integration [3], nonequilibrium work [4], MM/PBSA and MM/GBSA [5, 6], can offer reasonable accuracy, they are computational intensive and thus impractical for high-throughput applications like virtual screening, where scalability and robustness are essential.

In recent years, data-driven approaches have made significant progress and brought substantial promise to the field [7]. These methods can be broadly categorized by their input representations into sequence-based and structure-based models [8]. Sequence-based methods utilize only the protein amino acid sequences along with ligand information, without incorporating structural information of the proteins or complexes [9]. While such models avoid the need for structural data, they struggle to fully capture the intricate three-dimensional (3D) intermolecular interactions that fundamentally determine binding affinity, typically resulting in suboptimal performance and limited interpretability. In contrast, structure-based methods directly model the 3D structures of protein–ligand complexes, enabling the capture of geometric features and physical interactions between molecules, and offering potential insights into underlying binding mechanisms. These methods range from traditional machine learning models [10, 11], to 3D convolutional neural networks (CNNs) [12–14], and geometric graph neural networks (GNNs) [15, 16].

Despite these advances, the further development of structure-based affinity prediction has been significantly constrained by persistent data scarcity, stemming from the time-consuming and resource-intensive nature of experimentally determining protein–ligand complex structures.

For example, the PDBbind [17], the most widely used binding affinity benchmark dataset, contains fewer than 20, 000 complexes with annotated binding affinities. Moreover, many proteins and ligands within PDBbind exhibit high sequence or structural similarity, and this lack of diversity further limits the generalizability of models developed on the dataset.

To address this fundamental challenge, we investigate the potential of publicly available, large-scale, yet underexplored affinity measurements lacking corresponding experimental structural data, and develop a geometric deep learning–based scoring function, GatorAffinity, to advance affinity prediction. Current public databases, such as BindingDB [18], contain millions of protein–ligand interaction records with measured binding affinities; however, the vast majority lack corresponding 3D structures, limiting their utility for structure-based affinity model development. To bridge this gap, we construct GatorAffinity-DB, the first large-scale synthetic structural database centered on two of the most reliable affinity metrics: the dissociation constant (*K*_*d*_) and the inhibition constant (*K*_*i*_). We curate over 69, 000 high-quality *K*_*d*_ entries and 380, 000 *K*_*i*_ entries from BindingDB [18], and generate the missing complex structures using Boltz-1 [19], a leading structure prediction model. This process results in more than 450*K* newly synthesized structure–affinity pairs, expanding the scale of current structure-based datasets by a factor of 20. Additionally, we incorporated over 1 million *IC*_50_-labeled synthetic complexes from the recently developed SAIR database [20], which provides predicted 3D structures paired with affinity measurements. After structure correction and standardization, the final training dataset comprises over million synthetic structure–experimental affinity pairs, scaling existing resources by two orders of magnitude.

Leveraging this unprecedented scale of synthetic data, we develop GatorAffinity, a powerful geometric deep learning-based scoring function. Inspired by the recent work ATOMICA [21], which unsupervisedly learns atomic-scale representations of intermolecular interfaces across diverse biomolecular modalities, we adapt a similar GNN architecture specifically for protein–ligand affinity prediction. The model employs a hierarchical graph representation in which protein–ligand complexes are modeled at both the atom and block levels, with SE(3)-equivariant tensor field networks [22] performing message passing along intra- and inter-molecular edges. Building upon this architecture, GatorAffinity is pretrained on our large-scale synthetic structural dataset and subsequently fine-tuned on high-quality experimental data from PDBbind.

Extensive benchmark experiments demonstrate that our approach achieves significant performance gains, surpassing state-of-the-art (SOTA) structure-based and sequence-based protein–ligand affinity prediction methods. Additionally, we observe, for the first time, a data scaling law in affinity prediction: as the size of pre-training data increases, model performance improvements follow a power-law decay. This finding suggests that augmenting limited experimental data with large-scale, high-quality synthetic complexes can effectively overcome the data scarcity challenge while preserving strong predictive reliability—highlighting the immense potential of synthetic data to accelerate and transform drug discovery. By publicly releasing this synthetic structure–affinity database and associated scoring function, our work not only advances the protein-ligand binding affinity, but also unlocks new opportunities for scalable and accurate structure-based drug design. In summary, our main contributions are as follows:

- We construct GatorAffinity-DB, the largest synthetic structural dataset with annotated *K*_*d*_ and *K*_*i*_ binding affinities.
- We develop GatorAffinity, a powerful geometric deep learning–based scoring function, pretrained at scale and fine-tuned on high-quality experimental structures.
- Extensive experiments and ablation studies demonstrate that GatorAffinity achieves SOTA performance, highlighting the potential of synthetic structural data to significantly enhance binding affinity prediction.

## 2 Problem Formulation

Protein-ligand binding affinity prediction estimates the interaction strength between a protein *P* and a ligand *L*. In structure-based approaches, given a protein-ligand complex structure, the model predicts the binding affinity, typically measured by experimental metrics such as:

- *K*_*d*_ (Dissociation Constant): The equilibrium constant describing protein–ligand dissociation, directly reflecting the thermodynamic binding affinity. Values are obtained from direct binding assays.
- *K*_*i*_ (Inhibition Constant): The equilibrium dissociation constant for the binding of an inhibitor to an enzyme. Values depend on the kinetic mechanism of inhibition.
- *IC*_50_ (Half-maximal Inhibitory Concentration): Concentration reducing biological activity by 50% in a functional assay. Values highly depend on experimental conditions (e.g., substrate concentration) and inhibition mechanisms.

Among these metrics, *K*_*d*_ is considered the most reliable, *K*_*i*_ moderately reliable, and *IC*_50_ the least reliable. These values are converted to negative logarithmic scale *pK* (− log *K*_*d*_*/K*_*i*_*/IC*_50_) for model training and evaluation.

### Mathematical Formulation

A protein-ligand complex can be represented as a graph *G* = (*V*, ℰ), where *V* = *V*_*L*_ ∪ *V*_*P*_ .Here, *V*_*L*_ denotes the set of all ligand atoms, and *V*_*P*_ denotes the set of protein pocket atoms. The pocket is defined as the set of residues in which at least one atom lies within 5 Å of any ligand atom. Furthermore, ℰ= {ℰ_*ij*_ |*v*_*i*_, *v*_*j*_ ∈*V*} denotes the set of all edges constructed using *k*-NN. For each node *v*_*i*_ ∈ *V*, we extract its 3D coordinates **x**_*i*_ ∈ ℝ^3^ and atom features **a**_*i*_ ∈ ℝ^*m*^ (e.g., atom type). These are collected into an atom coordinate matrix **X** ∈ ℝ^*n×*3^ and an atom feature matrix **A** ∈ ℝ^*n×m*^, where *n* = |*V* | denotes the number of nodes in *G*. The edge features are denoted as 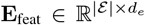, where each edge ℰ_*ij*_ has its own features (e.g. edge type). The binding affinity value *y* = *pK* ∈ ℝ is associated with the complex.

#### Input

A protein-ligand complex represented as graph *G* = (*V*, E).

#### Output

Predicted binding affinity *ŷ* ∈ ℝ.

#### Objective

Given a dataset 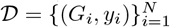, learn a function *f*_*θ*_ : *G* → ℝ that minimizes:

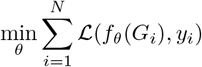

where ℒ is the loss function and *f*_*θ*_ is parameterized by *θ*.

## 3 GatorAffinity Database

Structural data of protein-ligand complexes with affinity annotations provide a crucial foundation for structure-based affinity model development. However, publicly available resources providing both structures and binding affinity remain scarce [26]. By contrast, affinity measurements without associated experimental structures are far more abundant. For instance, BindingDB [18] has accumulated millions of experimentally determined records, including *K*_*d*_, *K*_*i*_, and *IC*_50_ binding annotations.

Fortunately, with the rapid advancement of structure prediction models such as AlphaFold3 [27], Boltz-1 [19] and Chai-1 [28] in the past two years, the large-scale, high-precision computational generation of protein–small molecule complex structures has become a practical reality. This raises an important **question**: can these powerful models be leveraged to generate synthetic complex structure data that meaningfully enhance the development of structure-based affinity prediction methods by scaling the size of the training data? To investigate this large-scale yet underexplored data, we extracted reliable affinity measurements along with corresponding protein–ligand sequence information from BindingDB [18], and used Boltz-1 to generate structural conformations for these complexes—resulting in the GatorAffinity-DB dataset, with detailed statistics summarized in Table 1, alongside other publicly available protein–ligand structure–affinity databases.

**Table 1:** Public protein-ligand structure–affinity databases.

| Database | Structural Data Type | $K_d$ (entries) | $K_i$ (entries) | $IC_{50}$ (entries) |
| --- | --- | --- | --- | --- |
| PDBbind2013 [17] | Experimental | 4,682 | 3,028 | 3,066 |
| PDBbind2020 [17] | Experimental | 7,249 | 5,004 | 7,190 |
| Binding MOAD [23] | Experimental | 5,509 | 4,581 | 5,131 |
| BioLip 2 [24] | Experimental | 9,812 | 12,437 | 14,733 |
| HiQBind [25] | Experimental | 10,169 | 9,984 | 11,686 |
| SAIR [20] | Synthetic | N/A | N/A | **1,048,857** |
| **GatorAffinity-DB** | Synthetic | **69,201** | **387,325** | N/A |

### 3.1 Dataset Construction

To construct GatorAffinity-DB, we developed a comprehensive preprocessing workflow, beginning with the extraction of *K*_*d*_ and *K*_*i*_ measurements with their protein–ligand sequences from BindingDB. The multi-chain protein complexes and *K*_*i*_ measurements involving proteins with sequences longer than 1, 000 residues were excluded to reduce the computational costs and maintain structure prediction quality.

Additionally, since BindingDB aggregates binding affinity measurements from multiple literature sources, the same protein-ligand complex may have multiple affinity values reported from different experimental studies. To resolve these duplicate records, we retained the most frequently reported affinity value for each pair. In cases where multiple values shared the highest frequency, we used their arithmetic mean.

This approach ensures every unique complex has a single reliable affinity annotation while accounting for experimental variability across different literature sources.

Based on the filtered data, we applied Boltz-1 to predict the protein-ligand binding structure on the obtained data, yielding the 463, 867 complex samples with 70, 221 *K*_*d*_ and 393, 646 *K*_*i*_ entries. However, some of the Boltz-1-predicted structures exhibited physically invalid characteristics, including steric clashes between atoms, deviations in bond lengths and angles, incorrect stereochemistry at chiral centers, and so on.

To address these issues, we further filtered and corrected the predicted structures. Specifically, we employed the ligand-fixing pipeline from HiQBind [25], which includes modules for correcting bond orders and generating chemically reasonable protonation states. For protein structures, we used PDBFixer [29] to address missing atoms and structural inconsistencies. Following these filtering and structural correction, the final GatorAffinity-DB dataset comprises a total of 456, 526 complexes, including 69, 201 *K*_*d*_ and 387, 325 *K*_*i*_ entries.

### 3.2 Data Statistics

We analyzed the synthetic complexes in GatorAffinity-DB, examining distributions across protein sequence length, lig- and molecular weight, and affinity metrics (− log *K*_*d*_*/K*_*i*_). Furthermore, we evaluated ipTM scores (interface predicted TM-score) provided by Boltz-1, which reflect the prediction confidence of the protein–ligand interface. As shown in Fig. 1, the dataset demonstrates adequate diversity and quality. The ligand molecular weights concentrate within drug-like ranges, affinity values show physiologically relevant binding strengths, and most predictions have ipTM scores *>* 0.8, indicating high-confidence structural conformations.

**Figure 1:**
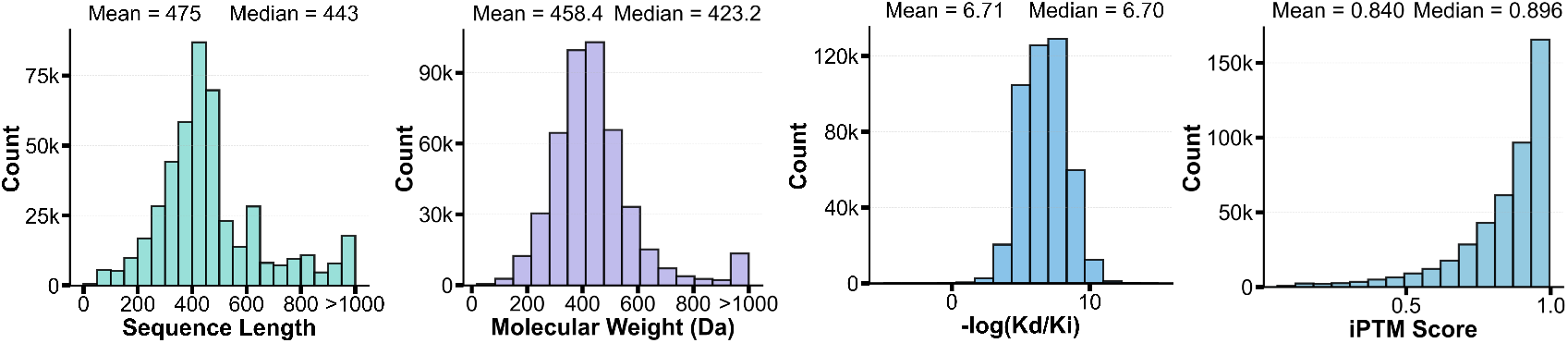
Statistical summary of the synthetic GatorAffinity-DB database.

## 4 GatorAffinity Framework

As illustrated in Fig. 2a-d, our complete GatorAffinity framework comprises four key stages, (1) sequence–affinity data collection, (2) synthetic structural data generation, (3) data processing and correction of synthetic structures, and (4) model development of the GatorAffinity scoring function. In this section, we first introduce our model architecture, followed by a description of the benchmark data preparation, and conclude with our model training strategy.

**Figure 2:**
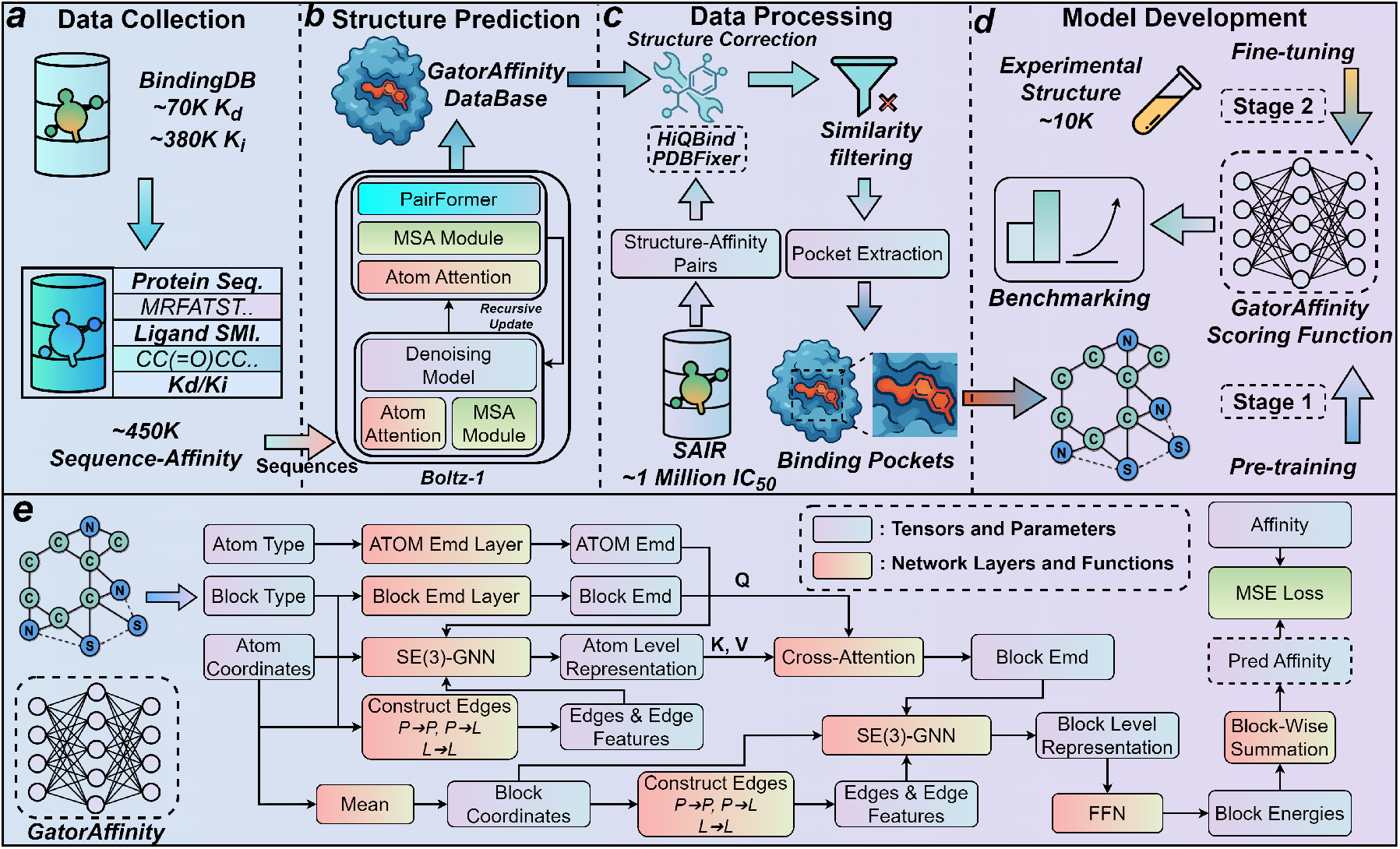
Overview of the GatorAffinity framework. **(a) Data Collection:** Binding affinity data sourced from BindingDB [18] (∼70K *K*_*d*_, ∼380K *K*_*i*_ measurements). **(b) Structure Prediction:** Protein-ligand complexes predicted using Boltz-1. **(c) Data Processing:** Structures corrected via HiQBind and PDBFixer, filtered for similarity, and binding pockets extracted. Combined with SAIR database (∼1M *IC*_50_ data) for pre-training. **(d) Model Development:** Model pre-trained on synthetic data, fine-tuned on experimental structures to reach SOTA. **(e) GatorAffinity Architecture:** Employs SE(3)-GNN layers, cross-attention, and hierarchical representations to predict binding affinity using block-wise energy computation.

### 4.1 Model Architecture

Our model extends ATOMICA’s hierarchical geometric deep learning framework for protein-ligand binding affinity prediction. As illustrated in Fig. 2e, each protein-ligand complex is represented as a hierarchical graph and the model generates multi-scale embeddings at both atomic and block levels.

#### Atom-level Graph

The complex is first decomposed into individual atoms, where each node corresponds to an atom with features including element type and 3D coordinates.

Edges are constructed based on spatial proximity, with intramolecular edges connecting each atom to its *k* nearest neighbors within the same molecule, and intermolecular edges connecting atoms across the protein-ligand interface.

#### Block-level Graph

Atoms are hierarchically grouped into chemically meaningful blocks. For proteins, these blocks correspond to residues from the 20 canonical amino acids, while for ligands, blocks represent functional moieties selected from a vocabulary of 290 common chemical fragments [21]. The coordinate of each block is defined as the mean position of its constituent atom coordinates.

#### SE(3)-Equivariant Message Passing

The model employs SE(3)-equivariant tensor field networks [22] for message passing, ensuring rotational and translational equivariance, strictly adhering to physical law and eliminating the need for data augmentation. At the atom level, the node representation is updated as follow:

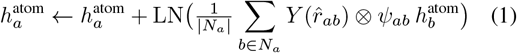

where 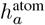 denotes the representation of atom *a*; *N*_*a*_ represents its neighbor set with size 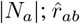 is the unit vector from atom *a* to atom *b*; *Y* (*·*) denotes the spherical harmonics; ⊗ indicates the tensor product; *ψ*_*ab*_ is a learned edge function derived from edge features; and LN(*·*) represents layer normalization.

The block-level representations are subsequently updated using a multi-head cross-attention mechanism between block nodes and their constituent atoms, enabling the model to aggregate fine-grained atomic information while preserving the hierarchical structure of the complex.

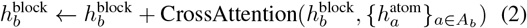

where 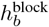 denotes the representation of block *b, A*_*b*_ represents the set of atoms associated with block *b*, and CrossAttention(,) represents the multi-head cross-attention mechanism.

#### Block-wise Energy Decomposition and Affinity Prediction

For binding affinity prediction, a specialized prediction head is first applied to each block-level representation 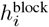 to compute its corresponding energy contribution 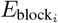 through a feed-forward network (FFN). Eventually, the energy contributions of all blocks within the protein–ligand complexes are then aggregated, by summation, to yield the final predicted binding affinity.

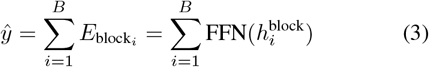

where *ŷ* represents the predicted affinity, *B* is the total number of blocks in the protein-ligand complex.

### 4.2 Benchmark Data Preparation

To fully explore the potential of synthetic structural data, we integrated our GatorAffinity-DB with the SAIR database, resulting in a training corpus of over 1.5 million synthetic structure-affinity pairs—representing a significant increase in scale compared to typical experimental datasets.

For evaluation, we used the LP-PDBbind dataset, a re-split version of PDBbind2020 dataset that mitigates data leakage through strict sequence similarity filtering. We further corrected numerous structural issues in LP-PDBBind—including incomplete ligands, incorrect bond annotations, coordinate displacements, and missing atoms—using the same refinement strategy applied to GatorAffinity-DB. Additionally, we removed entries containing buffer ligands, ligands with molecular weights exceeding 1, 000 Da, or irreparable structures to ensure that the benchmark set consisted of drug-like molecules with reliable structural quality. Since problematic or non–drug-like entries were primarily concentrated in the training set, we preserved the original data split while removing a total of 1, 717 entries, resulting in a filtered LP-PDBBind dataset, as summarized in Table 2. To prevent potential data leakage during the model pre-training, we followed the similarity filtering methodology utilized by LP-PDBBind and employed the same sequence similarity filtering between the integrated pretraining dataset and the experimental evaluation sets, thereby ensuring fair and reliable model evaluation.

**Table 2:** Size of LP-PDBBind before and after filtering.

| Subset | Pre-filtering | Post-filtering | Difference |
| --- | --- | --- | --- |
| Train | 11,513 | 10,212 | 1,301 |
| Validation | 2,422 | 2,201 | 221 |
| Test | 4,860 | 4,665 | 195 |
| Total | 18,795 | 17,078 | 1,717 |

### 4.3 Training Strategy

We propose a customized training strategy to fully exploit the scale of synthetic data while leveraging the accuracy of high-quality experimental data. The strategy consists of two stages: (1) pre-training on over 1.5 million synthetic structure–affinity pairs, followed by (2) fine-tuning on approximately 10, 000 experimental data points. This two-stage approach enables the model to first learn general protein–ligand interaction patterns and to derive a foundational representation of the relationship between binding modes and binding affinity using large-scale synthetic data. It is then refined on high-quality experimental structures to correct biases and mitigate distributional shifts introduced during synthetic pretraining. Both stages employ the mean squared error (MSE) loss function:

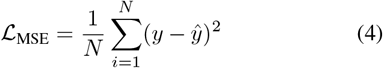

Meanwhile, prior to pretraining, we initialize our model with the ATOMICA weights, which were unsupervisedly pretrained on approximately 2 million biomolecular interaction complexes to capture fundamental knowledge of intermolecular interactions. Detailed ablation studies on different training strategies and model weight initialization are presented in Sections 5.4, 5.5, and 5.6.

## 5 Results

### 5.1 Benchmarking on the Filtered LP-PDBBind

We compared GatorAffinity with 12 existing models encompassing classical affinity prediction approaches and recent data-driven models. These include force field-based method such as AutoDock Vina [30], sequence-based deep learning methods [9, 31, 32], 3D structure-based CNN methods [33], and GNN-based methods [34–37, 15, 38, 16]. All baseline models were carefully re-trained on the same training set of the filtered LP-PDBBind dataset, with the validation set used to select optimal hyperparameters. The reported test results represent the mean and standard deviation across five independent runs. As shown in the Table 3, our GatorAffinity consistently outperforms all baseline approaches by a large margin across all evaluation metrics. Compared with the second-best model, GIGN—which is also structure-based—GatorAffinity achieves substantially lower root mean squared error (RMSE) and mean absolute error (MAE) by 11.0% and 11.2%, and higher Pearson’s *R* and Spearman *ρ*, correlations by 17.5% and 21.0%, respectively. When focusing on the therapeutically relevant affinity range of 1*mM* to 1*nM* [39], GatorAffinity consistently outperformed all baselines, with detailed results in the Table A.1. Moreover, GatorAffinity demonstrates superior performance on the benchmark datasets PDB-bind2016 and PDBbind2020, further confirming its robustness, as detailed in the Table A.2 and A.3.

**Table 3:** Performance comparison on the filtered LP-PDBBind dataset. Evaluation metrics include root mean squared error (RMSE) and mean absolute error (MAE) for prediction accuracy, Pearson’s correlation coefficient (*R*) and Spearman’s correlation coefficient (*ρ*) for correlation with experimental affinities, and the concordance index (CI) for ranking capability. The performance of all data-driven approaches is reported as mean ± standard deviation. The best results are highlighted with **bold text**.

| Model | RMSE (↓) | MAE (↓) | Pearson (↑) | Spearman (↑) | CI (↑) |
| --- | --- | --- | --- | --- | --- |
| AutoDock Vina | 2.285 | 1.709 | 0.365 | 0.383 | 0.634 |
| DeepDTA | 1.528 ± 0.025 | 1.215 ± 0.021 | 0.490 ± 0.023 | 0.452 ± 0.023 | 0.656 ± 0.008 |
| DrugBAN | 1.617 ± 0.028 | 1.276 ± 0.025 | 0.451 ± 0.016 | 0.447 ± 0.011 | 0.655 ± 0.004 |
| PSICHIC | 1.562 ± 0.031 | 1.227 ± 0.017 | 0.490 ± 0.018 | 0.472 ± 0.019 | 0.665 ± 0.006 |
| Pafnucy | 1.477 ± 0.068 | 1.181 ± 0.063 | 0.527 ± 0.235 | 0.502 ± 0.238 | 0.691 ± 0.085 |
| Interformer | 1.554 ± 0.035 | 1.228 ± 0.029 | 0.492 ± 0.030 | 0.459 ± 0.035 | 0.659 ± 0.012 |
| GraphDTA | 1.522 ± 0.007 | 1.205 ± 0.005 | 0.499 ± 0.012 | 0.481 ± 0.011 | 0.666 ± 0.004 |
| TANKBind | 1.534 ± 0.025 | 1.219 ± 0.022 | 0.490 ± 0.023 | 0.452 ± 0.022 | 0.656 ± 0.008 |
| TorchMD-Net | 1.531 ± 0.025 | 1.217 ± 0.021 | 0.490 ± 0.023 | 0.452 ± 0.023 | 0.656 ± 0.008 |
| SchNet | 1.529 ± 0.030 | 1.206 ± 0.020 | 0.506 ± 0.028 | 0.476 ± 0.031 | 0.665 ± 0.010 |
| EGNN | 1.476 ± 0.057 | 1.162 ± 0.042 | 0.542 ± 0.038 | 0.513 ± 0.038 | 0.680 ± 0.014 |
| GIGN | 1.453 ± 0.031 | 1.147 ± 0.019 | 0.572 ± 0.016 | 0.537 ± 0.012 | 0.689 ± 0.004 |
| **GatorAffinity** | **1.293 ± 0.005** | **1.018 ± 0.005** | **0.672 ± 0.003** | **0.650 ± 0.003** | **0.734 ± 0.001** |

### 5.2 Effectiveness of the Synthetic Data Pretraining

Our primary hypothesis is that incorporating synthetic structure–affinity data can enhance the development of structure-based affinity prediction methods by substantially increasing the scale of the training data. To validate it, we conducted experiments using varying amounts of synthetic data, including a setting without any synthetic data pretraining (as a baseline), and evaluated model performance under the same fine-tuning protocol. As shown in Fig. 3, the marginal gains in model performance follow a power-law decay as the size of the synthetic dataset increases, demonstrating that synthetic structural data can effectively enhance the generalization capability of structure-based affinity prediction models. Moreover, pre-training with ∼450*K Kd*+*Ki* data achieved results comparable to using ∼1*M IC*_50_ data, suggesting that both the quality of affinity measurements and the scale of data are critical factors in model development. The optimal performance was achieved only when all three data types were combined, indicating the importance of the sheer data quantity and diversity provided by the more abundant (albeit less precise) *IC*_50_ data.

**Figure 3:**
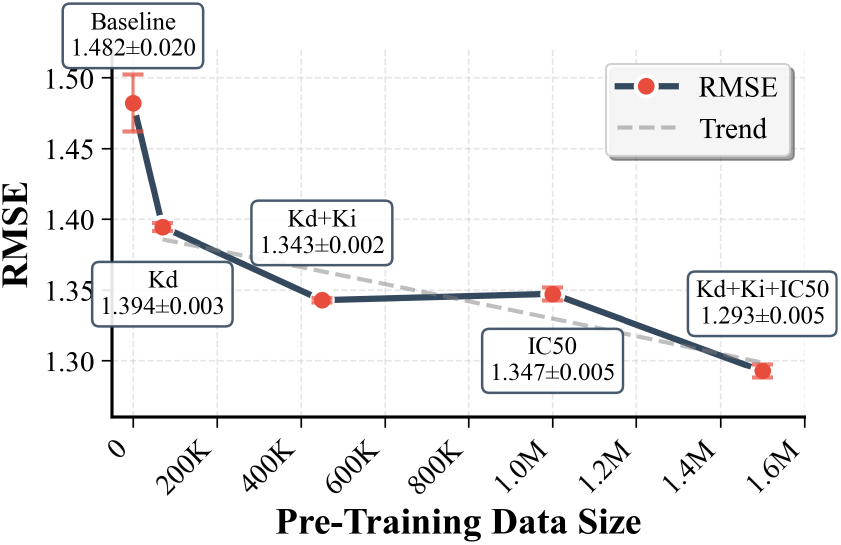
Relationship between pretraining dataset size and GatorAffinity RMSE on filtered LP-PDBBind

### 5.3 Ablation Study for ipTM Filtering

We investigated the impact of the structural quality of the synthetic data on model performance.

Previous methods typically apply strict ipTM filtering to retain only high-confidence structures to ensure data quality [20]. However, some studies suggest that incorporating noisier structures during training can, in certain cases, improve the model generalization capability [40].

Here, we applied an ipTM threshold of 0.75 to filter the synthetic data and retrained GatorAffinity using the resulting high-confidence subsets. The results indicate that, across all combinations of synthetic data, the impact of ipTM filtering on performance is minor and generally falls within the reported standard-deviation range, as shown by the comparison between the first and second blocks in Table 4. For example, the RMSE of the models pre-trained using *K*_*d*_ + *K*_*i*_ + *IC*_50_ data without ipTM filtering is 1.293± 0.005, compared with 1.300 ± 0.002 after applying the filtering. This suggests that the large-scale synthetic dataset strikes a balance between data quantity and structural noise, enabling robust final model performance without requiring aggressive filtering. Notably, although pretraining on low-confidence synthetic data may introduce noise, subsequent fine-tuning with high-quality experimental structures effectively mitigates these biases, leading models pre-trained with or without structural filtering strategies to converge to comparable performance. This outcome also highlights the effectiveness of experimental data fine-tuning (see Section 5.5) and two-stage training strategy (see Section 5.6).

**Table 4:** Ablation study of GatorAffinity on the filtered LP-PDBBind dataset.

| Variant | RMSE (↓) | MAE (↓) | Pearson (↑) | Spearman (↑) | CI (↑) |
| --- | --- | --- | --- | --- | --- |
| GatorAffinity (w/o pretraining) | 1.482 ± 0.020 | 1.173 ± 0.016 | 0.552 ± 0.012 | 0.538 ± 0.010 | 0.689 ± 0.004 |
| GatorAffinity ($K_d$) | 1.394 ± 0.003 | 1.110 ± 0.002 | 0.604 ± 0.002 | 0.577 ± 0.002 | 0.704 ± 0.001 |
| GatorAffinity ($K_d+K_i$) | 1.343 ± 0.002 | 1.062 ± 0.001 | 0.641 ± 0.001 | 0.619 ± 0.001 | 0.721 ± 0.000 |
| GatorAffinity ($IC_{50}$) | 1.347 ± 0.005 | 1.065 ± 0.003 | 0.637 ± 0.005 | 0.622 ± 0.004 | 0.722 ± 0.002 |
| GatorAffinity ($K_d+K_i+IC_{50}$) | 1.293 ± 0.005 | 1.018 ± 0.005 | 0.672 ± 0.003 | 0.650 ± 0.003 | 0.734 ± 0.001 |
| ipTM Filtering ($K_d$) | 1.403 ± 0.002 | 1.112 ± 0.002 | 0.598 ± 0.001 | 0.571 ± 0.001 | 0.702 ± 0.000 |
| ipTM Filtering ($K_d+K_i$) | 1.346 ± 0.011 | 1.064 ± 0.007 | 0.641 ± 0.004 | 0.616 ± 0.004 | 0.720 ± 0.002 |
| ipTM Filtering ($K_d+K_i+IC_{50}$) | 1.300 ± 0.002 | 1.020 ± 0.002 | 0.668 ± 0.001 | 0.648 ± 0.002 | 0.733 ± 0.001 |
| RandInit ($K_d$) | 1.492 ± 0.004 | 1.195 ± 0.004 | 0.530 ± 0.003 | 0.517 ± 0.003 | 0.680 ± 0.001 |
| RandInit ($K_d+K_i$) | 1.406 ± 0.002 | 1.115 ± 0.001 | 0.603 ± 0.002 | 0.577 ± 0.002 | 0.704 ± 0.001 |
| RandInit ($K_d+K_i+IC_{50}$) | 1.302 ± 0.003 | 1.026 ± 0.003 | 0.666 ± 0.001 | 0.643 ± 0.001 | 0.731 ± 0.001 |
| w/o ExpFineTune ($K_d$) | 1.563 ± 0.027 | 1.246 ± 0.011 | 0.514 ± 0.013 | 0.482 ± 0.012 | 0.668 ± 0.004 |
| w/o ExpFineTune ($K_d+K_i$) | 1.515 ± 0.003 | 1.213 ± 0.003 | 0.550 ± 0.002 | 0.527 ± 0.002 | 0.683 ± 0.001 |
| w/o ExpFineTune ($K_d+K_i+IC_{50}$) | 1.433 ± 0.005 | 1.147 ± 0.005 | 0.612 ± 0.002 | 0.597 ± 0.002 | 0.712 ± 0.001 |
| JointTrain ($K_d$) | 1.527 ± 0.003 | 1.210 ± 0.003 | 0.535 ± 0.007 | 0.517 ± 0.003 | 0.681 ± 0.001 |
| JointTrain ($K_d+K_i$) | 1.407 ± 0.006 | 1.114 ± 0.005 | 0.604 ± 0.004 | 0.579 ± 0.004 | 0.705 ± 0.002 |
| JointTrain ($K_d+K_i+IC_{50}$) | 1.377 ± 0.004 | 1.083 ± 0.004 | 0.622 ± 0.001 | 0.603 ± 0.001 | 0.715 ± 0.000 |

### 5.4 Ablation Study for Model Initialization

In this study, we initialized our model using parameters unsupervisedly pretrained by ATOMICA on large-scale biomolecular interaction data. This strategy allows GatorAffinity to begin training from a state that already encodes general knowledge of intermolecular interactions, rather than starting from random initialization. To assess the impact of this pretrained initialization, we conducted ablation experiments comparing model performance with and without pretrained weights.

As shown in Table 4, using random initialization (RandInit) leads to a certain degree of model performance decrease, but the magnitude of deterioration is inversely related to the amount of synthetic pretraining data. For example, when using only *K*_*d*_ data, RMSE increases from 1.394 to 1.492; in contrast, with *K*_*d*_ + *K*_*i*_ + *IC*_50_ data, the RMSE rises only from 1.293 to 1.302, leaving the overall performance virtually unaffected.

These results suggest a trade-off between data scale and model initialization: when training data is limited, prior knowledge from pretrained weights is crucial for performance; however, with sufficiently large-scale data (even if synthetic), the model can learn representations independently, reducing the relative contribution of pretrained initialization.

### 5.5 Effectiveness of Experimental Data Finetuning

To train GatorAffinity, we propose a two-stage strategy: large-scale pre-training using synthetic data, followed by fine-tuning on experimental structures. To evaluate the impact of this fine-tuning stage, we assessed models trained exclusively on synthetic data without subsequent refinement.

As shown in Table 4, removing the experimental fine-tuning stage (w/o ExpFineTune) leads to significant performance degradation across all synthetic data combinations. For instance, the model trained solely on synthetic *K*_*d*_ data, initialized with ATOMICA weights but without experimental fine-tuning, exhibits an RMSE increase from 1.394 to 1.563, accompanied by a Pearson correlation drop from 0.604 to 0.514. As the quantity of synthetic data increases, model performance steadily improves; however, a substantial gap remains compared with models fine-tuned on experimental structures (RMSE 1.433 vs. 1.293), even when the full set of approximately 1.5 million synthetic complexes is used. These results highlight the critical role of experimental structure fine-tuning in achieving optimal model performance, demonstrating that simply scaling the amount of synthetic training data is insufficient. The fidelity of synthetic structures continues to fall short of experimentally resolved complexes. This reinforces the urgent need for high-quality, large-scale, and openly available protein–ligand datasets to advance binding affinity model development, aligning with the recent initiative by Edwards et al. [41].

### 5.6 Ablation Study for Our Two-stage Training

To further demonstrate the two-stage training strategy’s effectiveness, here we compared it with an alternative one-stage setting. In this setting, joint training was performed on both synthetic and experimental data simultaneously, with weighted sampling applied to ensure that each training batch contained 20% experimental data. As shown in Table 4, the joint training strategy (JointTrain) consistently underperforms our two-stage approach across all data combinations. For *K*_*d*_ +*K*_*i*_ +*IC*_50_, one-stage training achieves an RMSE of 1.377 compared to 1.293 for our method. The performance gap is particularly pronounced for smaller datasets: with *K*_*d*_-only data, one-stage training yields an RMSE of 1.527 versus 1.394. These results indicate that simply mixing synthetic and experimental data leads to inadequate performance. This limitation likely arises from two key challenges: (1) the model treats synthetic and experimental data equivalently despite substantial differences in quality, and (2) distributional mismatches between the two data sources can introduce prediction biases.

By contrast, our two-stage strategy explicitly distinguishes between high- and low-fidelity data, enabling the model to first learn general interaction patterns from large-scale synthetic data and then adapt to small-scale, high-quality experimental structures for refinement.

## 6 Conclusion

In this study, we addressed the long-standing challenge of data scarcity in structure-based protein-ligand binding affinity prediction. We introduced GatorAffinity-DB, the first large-scale synthetic structural database focused on experimental *K*_*d*_ and *K*_*i*_ measurements, containing over 450, 000 predicted protein-ligand complexes. Leveraging GatorAffinity-DB together with the SAIR dataset, we developed GatorAffinity, a hierarchical geometric deep learning model-based scoring function, trained with a two-stage strategy: large-scale pre-training on synthetic structures followed by experimental data fine-tuning. Extensive experiments and ablation studies show that GatorAffinity consistently and substantially outperforms existing SOTA methods.

To our knowledge, this is the first work to demonstrate the benefits of synthetic structural data for protein–ligand binding affinity prediction and to reveal a power-law relationship between dataset size and model performance. Extrapolating from this relationship, systematic dataset expansion, augmented by the rapid evolution of structure prediction tools, promises significant and predictable improvements in model performance. Combined with our released synthetic datasets and pretrained models, these advances provide actionable pathways for accelerating structure-based drug discovery.

## Code and Data Availability

The source code for GatorAffinity, and the corresponding model checkpoints, and GatorAffinity-DB are publicly available via GitHub at https://github.com/AIDD-LiLab/GatorAffinity. The GatorAffinity-DB dataset can be accessed at https://huggingface.co/datasets/AIDD-LiLab/GatorAffinity-DB.

## Acknowledgement

This work was supported in part by the University of Florida (UF Startup Fund, UF Health Cancer Center Pilot Grant # UFS-2023-08, and UF Research AI Award to Y. L.), National Institutes of Health (R21EB037868 to Y. L.), and the Bodor Professorship Fund (to C. L.). We acknowledge UFIT Research Computing for providing computational resources.

## Appendix A Additional Experimental Results

### A.1 Benchmarking on the Drug-Relevant Affinity Range (1*mM* − 1*nM* )

To evaluate model performance within the therapeutically relevant affinity range, we filtered the LP-PDBbind dataset, spanning training, validation, and test sets, to include only complexes with binding affinities between 1*mM* and 1*nM*. This range is particularly important for drug discovery, as it encompasses the vast majority of successful pharmaceutical compounds [1] while excluding extreme affinity values that are often affected by experimental uncertainties or limited clinical relevance [2].

We then evaluated GatorAffinity and all baseline models in this focused dataset, which retained approximately 89% of the original set of complexes. As shown in Table A.1, while all models exhibited performance improvements on this focused dataset, GatorAffinity consistently achieved the best performance within this pharmaceutically relevant range.

### A.2 Benchmarking on the PDBbind2020

To further assess the scoring power of our approach, we evaluated GatorAffinity on the widely-used PDBbind2020 benchmark [3]. As shown in Table A.2, GatorAffinity achieved an RMSE of 1.281 ± 0.029 (mean ± SD), demonstrating clear improvements over state-of-the-art methods, such as, PSICHIC (1.328 ± 0.069) and GIGN (1.371 ± 0.054). The advantage is also evident in correlation metrics, where our model attained a Pearson correlation coefficient of 0.733 ± 0.012 (mean ± SD), compared with 0.720 ± 0.011 for PSICHIC and 0.680 ± 0.029 for GIGN. These results further validate the superior performance of our proposed GatorAffinity scoring function.

### A.3 Benchmarking on the PDBbind2016

We also validated our approach on PDBbind2016 [3] bench-marking dataset. GatorAffinity maintains its state-of-the-art performance with an RMSE of 1.279 ± 0.007 (mean ± SD), outperforming all the baseline approaches, e.g., GIGN (1.379 ± 0.047) and PSICHIC (1.356 ± 0.032). The consistent improvement across different dataset versions demonstrates the robustness of our method.

It is important to acknowledge that both PDBbind benchmarks have known limitations due to high sequence similarity between training and test proteins, which can lead to data leakage and inflated performance metrics. This issue affects all models evaluated on these benchmarks. Despite these limitations, our substantial improvements suggest that synthetic data pre-training provides genuine benefits beyond potential memorization effects. Moreover, the consistent advantages of GatorAffinity across multiple benchmarks, including the more stringent LP-PDBbind evaluation, further validate the effectiveness and robustness of our approach.

## B Additional Details on GatorAffinity-DB

### B.1 Statistics of *K*_*d*_ and *K*_*i*_ Subsets in GatorAffinity-DB

Fig. B.1 presents the independent statistical distributions of *K*_*d*_ (top panel) and *K*_*i*_ (bottom panel) entries in GatorAffinity-DB. The clear difference in protein sequence length distributions between two subsets arises from the inclusion of proteins with sequence lengths greater than 1, 000 residues in the *K*_*d*_ set. Furthermore, the − log(*K*_*d*_*/K*_*i*_) values also differ between the subsets, with the *K*_*d*_ entries having a median of 5.68 and the *K*_*i*_ entries have a median of 6.80. Note that both subsets maintain high structural prediction confidence, with the majority of data having ipTM scores above 0.8.

### B.2 Synthetic Structure Fix

To date, although state-of-the-art structure prediction models have demonstrated strong overall performance, some predicted structures still exhibited physically invalid characteristics, including steric clashes between atoms, deviations in bond lengths and angles, incorrect stereochemistry at chiral centers, and stereogenic bonds, non-planar aromatic rings, and ligands with improperly annotated bond types—often labeled uniformly as single bonds, lacking accurate bond orders and connectivity information. Such deficiencies introduce noise into synthetic structural data, pose challenges for parsing by cheminformatics toolkits such as RDKit [4], and potentially impede the development of reliable affinity prediction models. To address these issues, we used HiQbind [5] and PDBFixer [6] to fix all ligands and proteins, respectively.

## C GatorAffinity Training Details

### C.1 Implementation Environment

All experiments were conducted using PyTorch 2.7.0 with CUDA 12.8 support as the deep learning framework. The PyTorch ecosystem included PyTorch Geometric 2.6.1 for graph neural network operations, along with supporting libraries torch-cluster 1.6.3 and torch-scatter 2.1.2 for efficient graph computations. Pre-training was performed on 4 NVIDIA B200 GPUs, while fine-tuning experiments were carried out on 3 NVIDIA L4 GPUs.

### C.2 Pre-training Hyperparameters

We initialized our base model for atom and block representation learning using the pre-trained ATOMICA [7] foundation model, configured with: atom hidden size of 32, block hidden size of 32, edge size of 32, and 4 layers. The model employed *k*-nearest neighbors graph construction with *k* = 8, applied separately for intra- and inter-molecular edges, and a ligand fragmentation method based on a vocabulary of 290 common chemical fragments, where any unassigned atoms form their own single-atom blocks [7]. The model type was `DenoisePretrainModel` with 436 masked block classes [7].

**Table A.1:** Performance of each model on focused LP-PDBbind (1 mM − 1 nM). The best results are highlighted with **bold text**.

| Model | RMSE (↓) | MAE (↓) | Pearson (↑) | Spearman (↑) | CI (↑) |
| --- | --- | --- | --- | --- | --- |
| AutoDock Vina | 2.115 | 1.595 | 0.335 | 0.353 | 0.622 |
| DeepDTA | $1.347 \pm 0.031$ | $1.074 \pm 0.025$ | $0.425 \pm 0.021$ | $0.401 \pm 0.022$ | $0.637 \pm 0.008$ |
| DrugBAN | $1.385 \pm 0.026$ | $1.113 \pm 0.025$ | $0.399 \pm 0.018$ | $0.390 \pm 0.016$ | $0.634 \pm 0.006$ |
| PSICHIC | $1.278 \pm 0.015$ | $1.024 \pm 0.009$ | $0.490 \pm 0.010$ | $0.467 \pm 0.008$ | $0.661 \pm 0.003$ |
| Pafnucy | $1.375 \pm 0.003$ | $1.128 \pm 0.006$ | $0.456 \pm 0.001$ | $0.450 \pm 0.003$ | $0.658 \pm 0.002$ |
| Interformer | $1.251 \pm 0.015$ | $1.006 \pm 0.007$ | $0.496 \pm 0.010$ | $0.468 \pm 0.005$ | $0.662 \pm 0.002$ |
| GraphDTA | $1.288 \pm 0.015$ | $1.046 \pm 0.008$ | $0.448 \pm 0.012$ | $0.440 \pm 0.007$ | $0.651 \pm 0.003$ |
| TANKBind | $1.265 \pm 0.036$ | $1.014 \pm 0.025$ | $0.491 \pm 0.041$ | $0.475 \pm 0.026$ | $0.665 \pm 0.009$ |
| TorchMD-Net | $1.330 \pm 0.026$ | $1.061 \pm 0.015$ | $0.442 \pm 0.004$ | $0.417 \pm 0.006$ | $0.643 \pm 0.002$ |
| SchNet | $1.246 \pm 0.021$ | $1.004 \pm 0.007$ | $0.501 \pm 0.015$ | $0.473 \pm 0.001$ | $0.663 \pm 0.000$ |
| EGNN | $1.262 \pm 0.037$ | $1.013 \pm 0.026$ | $0.491 \pm 0.041$ | $0.475 \pm 0.026$ | $0.665 \pm 0.009$ |
| GIGN | $1.249 \pm 0.018$ | $1.001 \pm 0.017$ | $0.502 \pm 0.023$ | $0.483 \pm 0.011$ | $0.668 \pm 0.004$ |
| **GatorAffinity** | **$1.134 \pm 0.003$** | **$0.906 \pm 0.004$** | **$0.613 \pm 0.002$** | **$0.597 \pm 0.002$** | **$0.712 \pm 0.001$** |

**Table A.2:** Performance comparison on the PDBbind 2020. The best results are highlighted with **bold text**.

| Model | RMSE (↓) | MAE (↓) | Pearson (↑) | Spearman (↑) | CI (↑) |
| --- | --- | --- | --- | --- | --- |
| AutoDock Vina | 2.568 | 1.894 | 0.339 | 0.375 | 0.624 |
| DeepDTA | $1.468 \pm 0.047$ | $1.173 \pm 0.036$ | $0.644 \pm 0.016$ | $0.611 \pm 0.012$ | $0.718 \pm 0.007$ |
| DrugBAN | $1.471 \pm 0.041$ | $1.141 \pm 0.040$ | $0.634 \pm 0.021$ | $0.586 \pm 0.025$ | $0.711 \pm 0.010$ |
| PSICHIC | $1.328 \pm 0.069$ | $1.043 \pm 0.066$ | $0.720 \pm 0.011$ | $0.693 \pm 0.005$ | $0.753 \pm 0.003$ |
| Pafnucy | $1.444 \pm 0.013$ | $1.150 \pm 0.011$ | $0.612 \pm 0.011$ | $0.569 \pm 0.010$ | $0.760 \pm 0.001$ |
| Interformer | $1.374 \pm 0.058$ | $1.099 \pm 0.036$ | $0.688 \pm 0.023$ | $0.636 \pm 0.026$ | $0.729 \pm 0.011$ |
| GraphDTA | $1.514 \pm 0.034$ | $1.190 \pm 0.020$ | $0.584 \pm 0.018$ | $0.527 \pm 0.028$ | $0.686 \pm 0.011$ |
| TANKBind | $1.370 \pm 0.061$ | $1.096 \pm 0.039$ | $0.688 \pm 0.025$ | $0.639 \pm 0.027$ | $0.730 \pm 0.011$ |
| TorchMD-Net | $1.388 \pm 0.021$ | $1.100 \pm 0.020$ | $0.683 \pm 0.016$ | $0.639 \pm 0.024$ | $0.731 \pm 0.009$ |
| SchNet | $1.373 \pm 0.064$ | $1.100 \pm 0.039$ | $0.689 \pm 0.025$ | $0.639 \pm 0.027$ | $0.730 \pm 0.011$ |
| EGNN | $1.387 \pm 0.020$ | $1.100 \pm 0.018$ | $0.683 \pm 0.016$ | $0.639 \pm 0.024$ | $0.730 \pm 0.009$ |
| GIGN | $1.371 \pm 0.054$ | $1.084 \pm 0.050$ | $0.680 \pm 0.029$ | $0.635 \pm 0.038$ | $0.729 \pm 0.015$ |
| **GatorAffinity** | **$1.281 \pm 0.029$** | **$0.995 \pm 0.023$** | **$0.733 \pm 0.012$** | **$0.719 \pm 0.018$** | **$0.765 \pm 0.008$** |

**Table A.3:** Performance Comparison on the PDBbind2016. The best results are highlighted with **bold text**.

| Model | RMSE (↓) | MAE (↓) | Pearson (↑) | Spearman (↑) | CI (↑) |
| --- | --- | --- | --- | --- | --- |
| AutoDock Vina | 1.843 | 1.451 | 0.615 | 0.614 | 0.719 |
| DeepDTA | 1.473 ± 0.022 | 1.178 ± 0.033 | 0.746 ± 0.012 | 0.737 ± 0.011 | 0.772 ± 0.005 |
| DrugBAN | 1.546 ± 0.022 | 1.215 ± 0.018 | 0.727 ± 0.005 | 0.769 ± 0.003 | 0.494 ± 0.014 |
| PSICHIC | 1.356 ± 0.032 | 1.045 ± 0.026 | 0.784 ± 0.012 | 0.771 ± 0.017 | 0.793 ± 0.007 |
| Pafnucy | 1.679 ± 0.149 | 1.355 ± 0.119 | 0.675 ± 0.044 | 0.692 ± 0.060 | 0.749 ± 0.023 |
| Interformer | 1.500 ± 0.034 | 1.195 ± 0.010 | 0.729 ± 0.016 | 0.727 ± 0.017 | 0.766 ± 0.008 |
| GraphDTA | 1.622 ± 0.025 | 1.292 ± 0.022 | 0.686 ± 0.009 | 0.671 ± 0.010 | 0.744 ± 0.003 |
| TANKBind | 1.401 ± 0.061 | 1.123 ± 0.035 | 0.774 ± 0.022 | 0.769 ± 0.024 | 0.787 ± 0.010 |
| TorchMD-Net | 1.403 ± 0.061 | 1.126 ± 0.035 | 0.774 ± 0.022 | 0.769 ± 0.023 | 0.786 ± 0.010 |
| SchNet | 1.498 ± 0.022 | 1.195 ± 0.012 | 0.730 ± 0.014 | 0.729 ± 0.015 | 0.768 ± 0.007 |
| EGNN | 1.408 ± 0.062 | 1.129 ± 0.036 | 0.774 ± 0.022 | 0.769 ± 0.024 | 0.786 ± 0.010 |
| GIGN | 1.379 ± 0.047 | 1.096 ± 0.047 | 0.782 ± 0.014 | 0.776 ± 0.017 | 0.791 ± 0.007 |
| **GatorAffinity** | **1.279 ± 0.007** | **1.014 ± 0.008** | **0.813 ± 0.004** | **0.803 ± 0.005** | **0.807 ± 0.003** |

**Figure B.1:**
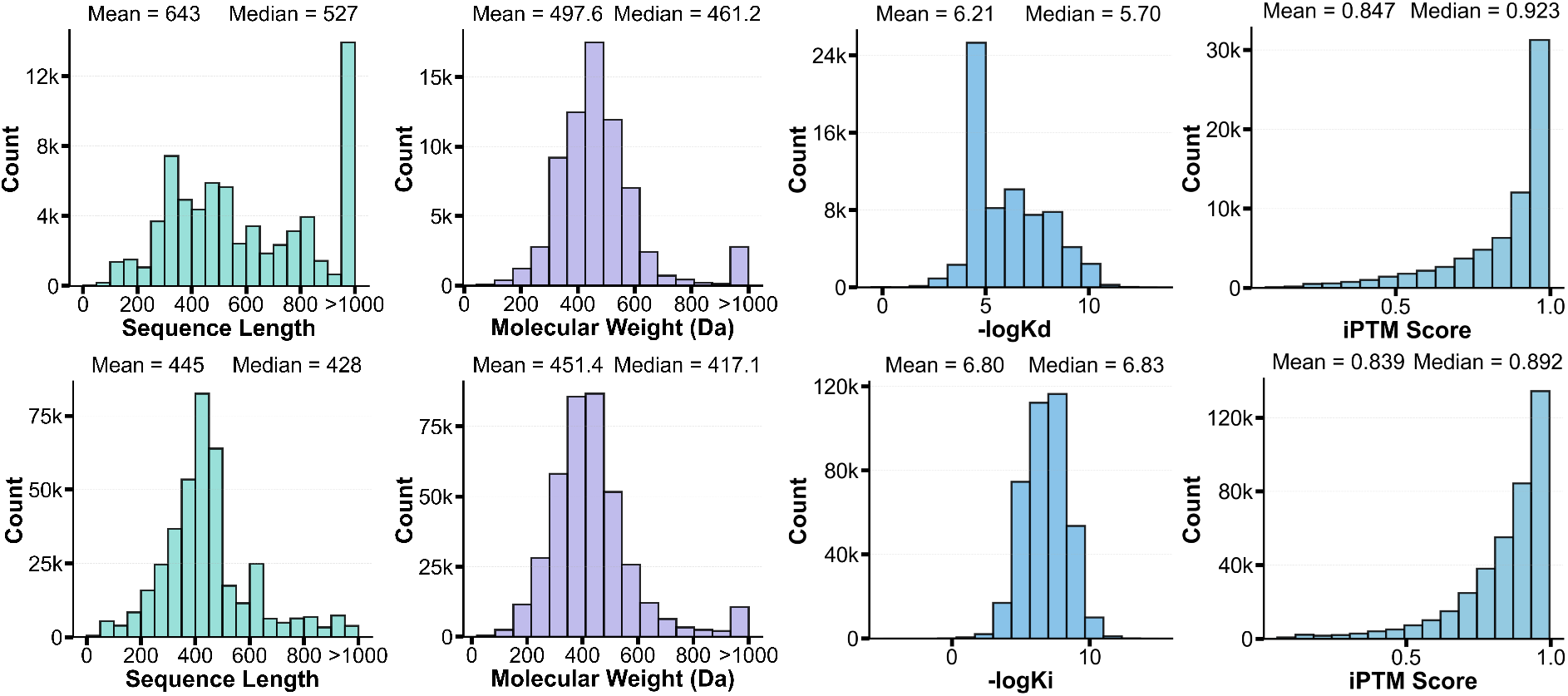
Statistical summary of Kd and Ki values in the synthetic GatorAffinity-DB

For the binding affinity prediction head, we implemented a 3-layer multilayer perceptron (MLP) with architecture [32, 32] → [32, 32] → [32, 1], where ReLU activation functions were applied between layers.

The first stage of synthetic data pre-training was conducted in a batch size of 450, with an initial learning rate of 1 × 10^−5^. A learning rate scheduling was implemented using cosine annealing with warm restarts, specifically CosineAnnealingWarmRestarts with *T*_0_ = 5, *T*_*mult*_ = 2, and *η*_*min*_ = 1 × 10^−6^. The random seed was set to 1 for reproducibility. No additional dropout regularization was applied (dropout rate = 0).

### C.3 Fine-tuning Hyperparameters

During fine-tuning, the learning rate was reduced to 1 × 10^−6^ with a smaller batch size of 55. The cosine annealing scheduler maintained the same configuration (*T*_0_ = 5, *T*_*mult*_ = 2) but with a lower minimum learning rate of *η*_*min*_ = 1 × 10^−7^. Random seeds from 1 to 5 were assigned to ensure experimental reproducibility and to facilitate statistical evaluation across five independent runs. All other architectural parameters, including the MLP configuration and model architecture, remained identical to the pre-training phase.

## D Pseudocode for GatorAffinity

We summarize our framework in Algorithm 1 and 2.

### Algorithm 1

Training Process for GatorAffinity

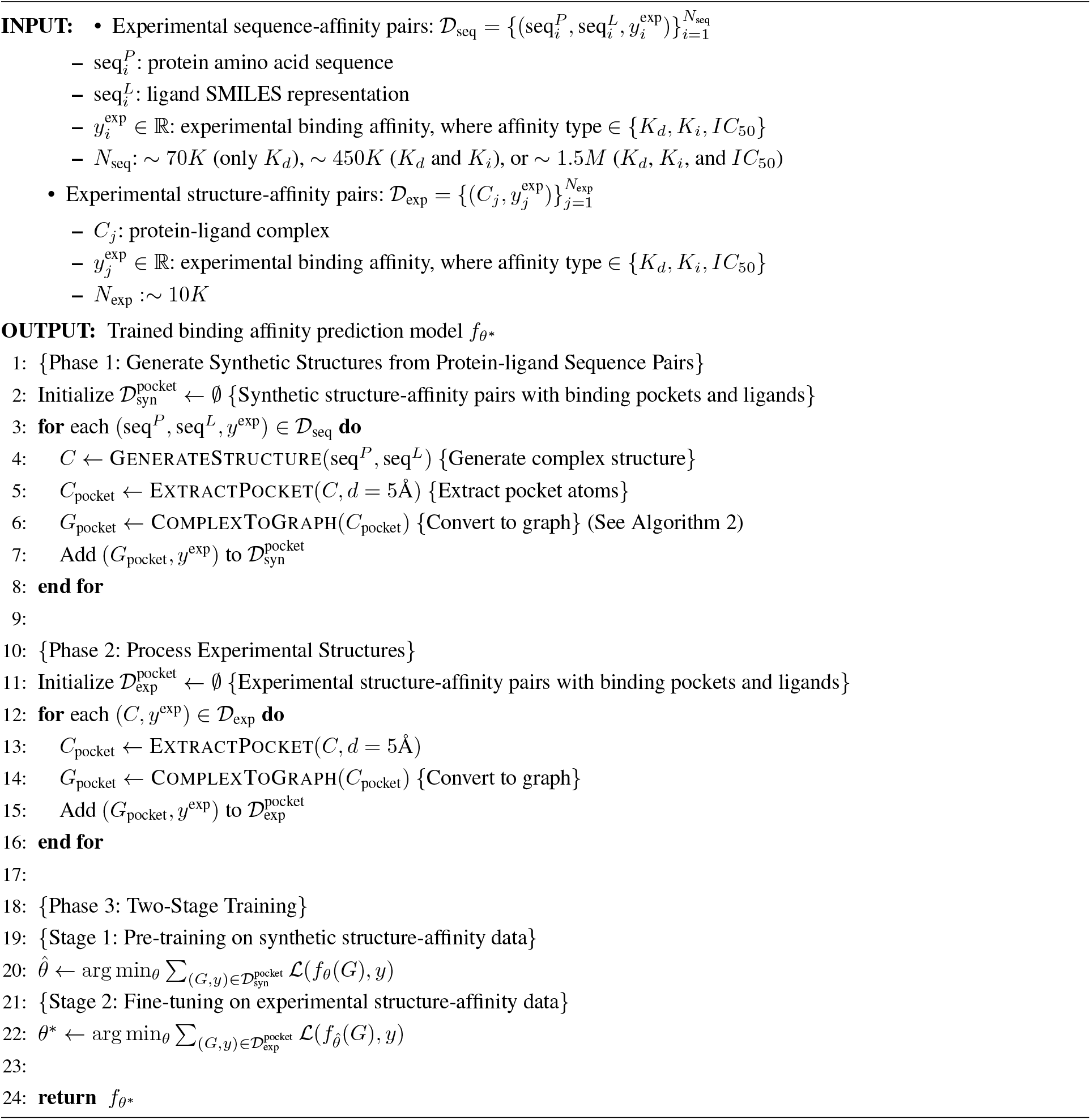

### Algorithm 2

ComplexToGraph: Converting Protein-Ligand Complex to Graph

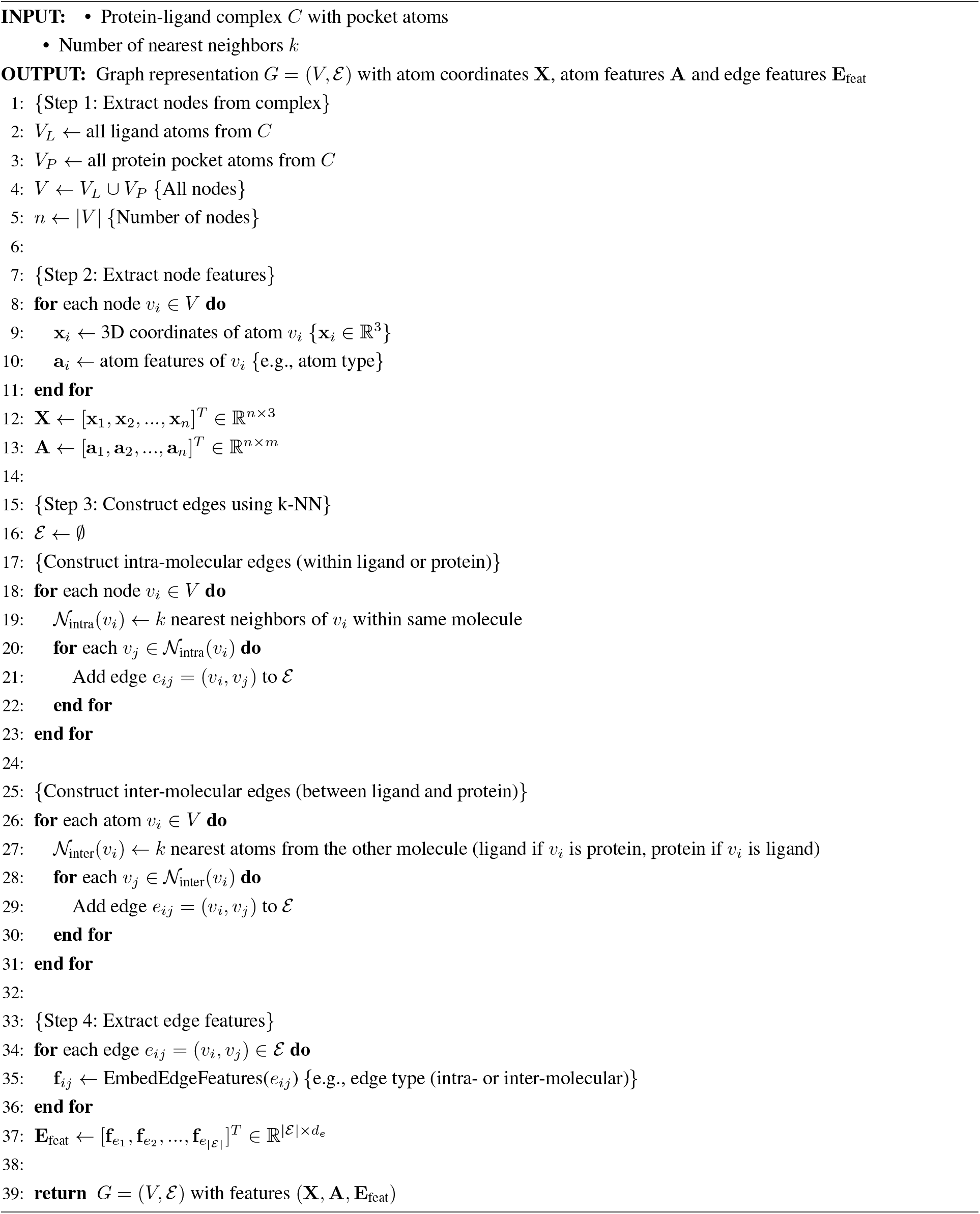

